# Key role of Pro230 in the hinge region on the IgG architecture and function

**DOI:** 10.1101/2024.05.10.593077

**Authors:** Yuuki Koseki, Yuki Yamaguchi, Michihiko Aoyama, Minoru Tada, Akinobu Senoo, Akiko Ishii-Watabe, Takayuki Uchihashi, Susumu Uchiyama, Koichi Kato, Saeko Yanaka, Jose M.M. Caaveiro

**Author notes:** Corresponding authors: Saeko Yanaka, Jose M.M. Caaveiro.

## Abstract

Immunoglobulin G (IgG) is a molecule that plays an important role in biological defense; IgG molecules have been applied as drugs due to their high specificity for antigens and their ability to activate immunity via effector molecules on immune cells. On the other hand, the flexibility of the hinge region makes it difficult to apply conventional structural biology approaches due to its dynamic conformational changes, and the mechanism of action of the molecule as a whole has not been elucidated. Here, we introduced a deletion amino acid mutation in the hinge region to elucidate the role of the hinge region and its effect on the structure and function of the IgG molecule. Deletion of Pro230 resulted in the formation of a half-molecular in which the interaction between heavy chains was lost. We elucidated the mechanism of half-IgG formation by structural analysis using nuclear magnetic resonance (NMR) measurements and by disulfide quantification using peptide mapping using LC-MS/MS. For this purpose, a new NMR stable isotope labeling method was introduced. Finally, cell assay revealed that the IgG half-molecules have specific FcγRI-mediated activity. This report provides new insights into the higher-order structure formation of IgG molecules and is expected to contribute to the elucidation of the molecular basis of the Fcγ receptor-mediated activation mechanism of the immune system.

## Introduction

Immunoglobulin G (IgG) is a major molecule working in biological defense in the human immune system. IgG consists of two identical light chains consisting of V_L_ and C_L_ domains and two identical heavy chains consisting of V_H_, C_H_1, C_H_2 and C_H_3 domains, with the C_H_1 and C_H_2 domains linked by a flexible hinge region. The two heavy chains form a homodimer due to two pairs of disulfide bonds in the hinge region and interactions between C_H_3 domains.^1^ The IgG structure can be also explained by their functional regions; the Fab region that recognizes antigen molecule, the Fc region that binds to effector molecule, and the hinge region that connects them. The flexibility of the hinge region makes it difficult to apply conventional structural biology approaches, such as X-ray crystallography, to the IgG undergoing dynamic conformational changes in solution.^2–4^ Therefore, the behavior of the molecular as a whole in vivo and its molecular mechanisms of action remain unclear. Especially, the hinge region is thought to link the functions of the Fab and Fc regions, and previous studies have reported the effect of the upper hinge and core hinge region on the effector function of IgG1.^5–14^ However, the detailed role of the hinge region at molecular level remains to be unclear.

IgG exerts its immune function by mediating highly specific antigen binding in Fab and its ability to activate the immune system by binding to effector molecules such as Fcγ receptors. Fcγ receptors are a family of proteins expressed on effector cells, including NK cells and macrophages. There are five subtypes of human Fcγ receptors (FcγRs): FcγRI, RIIa, RIIb, RIIIa and RIIIb. Each receptor is involved in the regulation of effector functions such as B cell activation, dendritic cell maturation, phagocytosis, antibody-dependent cellular cytotoxicity (ADCC), and inflammatory cell recruitment.^15–17^

While some FcγR such as FcγRIIIa has been studied extensively^18–23^, FcγRI (CD64) is the least known among the FcγRs. FcγRI is a glycoprotein expressed on macrophages and monocytes and is reported to contribute to phagocytic activity in the immune system by binding to the IgG molecule^24–27^. FcγRI is the only high affinity receptor in the Fcγ family. The FcγRI comprises three extracellular domains (termed D1, D2 and D3), a transmembrane domain, and an intracellular tail. The binding mode of the FcγRI and Fc is well known by the crystal structure of the complex of FcγRI and Fc. The structure indicates that the interaction between Fc and the receptor involves domains D1 and D2 of FcγRI, and the C_H_2 domain and hinge region of Fc.^15,28,29^

To elucidate the role of the hinge region, and to examine the effect of mutation on the structure and function of the IgG molecule, we conducted structural analysis and cell-based assays on a series of lower hinge mutations of Trastuzumab, a humanized antibody of the IgG1 class. Deletion of the residue Pro230 resulted in the formation of an IgG half-molecule, in which the interactions between IgG heavy chains are lost. We elucidated the mechanism of half-IgG formation by structural analysis using nuclear magnetic resonance (NMR) measurements and disulfide quantification using peptide mapping by LC-MS/MS. Interestingly, cell-based assays also revealed that the resulting half IgG still displays effector activity mediated by FcγRI, but not appreciably by other FcγRs.

## Results

### Mutagenesis in the low hinge of IgG1 generates half-antibody moieties

Deletion mutants in the lower hinge region of Trastuzumab, a humanized antibody of the IgG1 family, were systematically introduced to understand the function of this region on the conformational stability of this class of antibodies. Specifically, residues Pro230∼Leu234 were modified one by one as follows: ΔP230, ΔA231, ΔP232, ΔE233, ΔL234. The double substitution Cys226Ser and Cys229Ser termed CSCS was also investigated (Fig, 1a). The unmodified antibody (WT) and its mutants were expressed in a mammalian cell expression system, followed by their purification to homogeneity as described in the Materials and Methods section.

Analysis of WT and mutants by SDS-PAGE under reducing conditions showed a band corresponding to the heavy chain at around 50 kDa and a lower molecular weight band corresponding to the light chain at around 25 kDa (Fig. 1b). In contrast, under non-reducing conditions the SDS-PAGE revealed two different patterns. For WT and several mutants, a band at around 150 kDa were observed, corresponding to the fully assembled dimeric IgG1. As expected, the CSCS double mutant lacking disulfide bonds in the hinge region appeared at around 75 kDa, corresponding to a half-sized antibody. Surprisingly, the bands corresponding to the deletion mutant ΔP230 appeared as a major fraction at a position of 75 kDa (half-size antibody) and a minor fraction consistent with the full-size antibody at 150 kDa (Fig.1b). This result indicated that, to a large extent, disulfide bonds between the heavy chains in the lower hinge of deletion mutant ΔP230 are abrogated.

**Figure 1.**
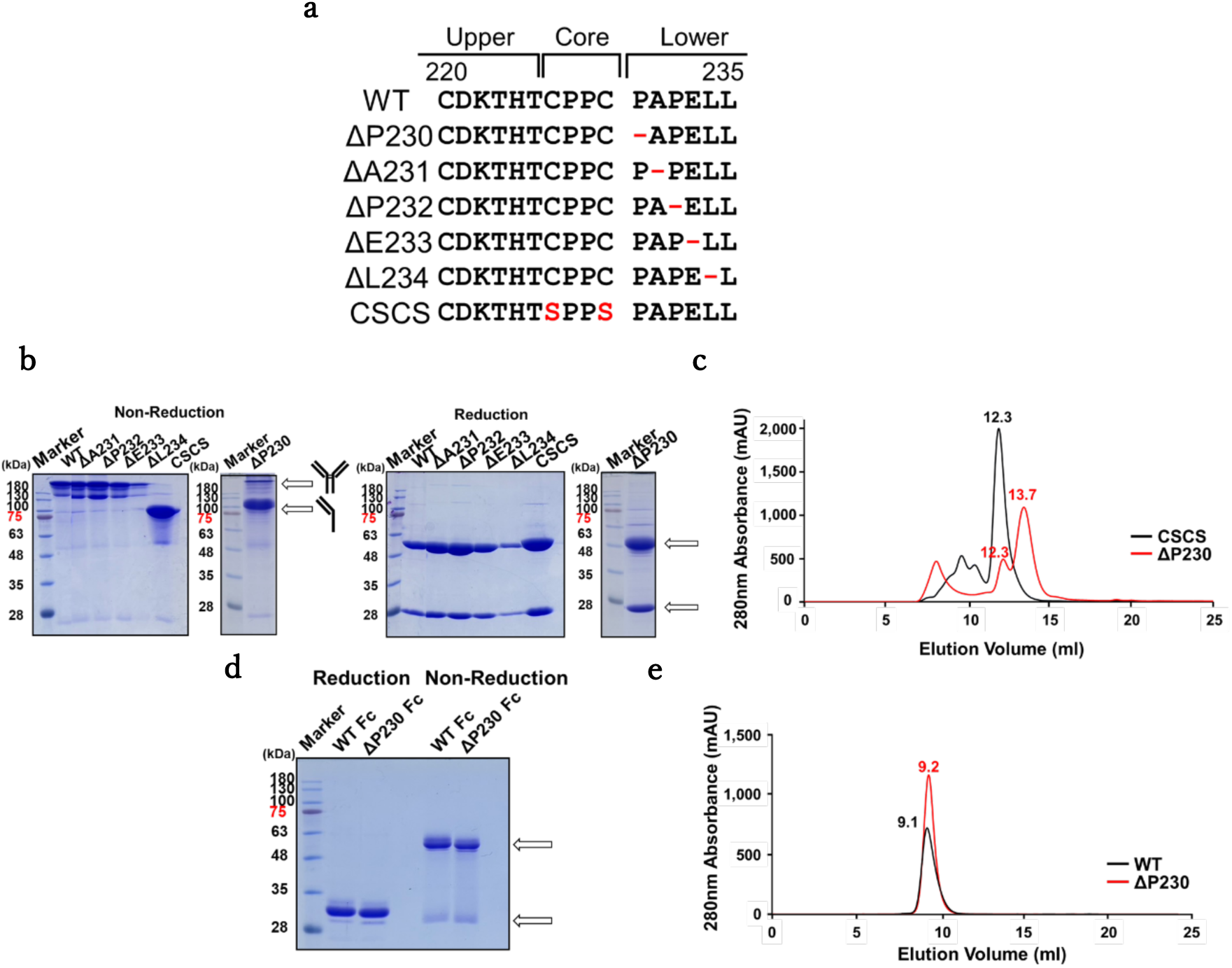
Preparation of lower hinge mutants of full length IgG1 (trastuzumab) and its Fc region. **(a)** Primary sequence of the lower hinge of human IgG1 (Cys220-Leu235) and the mutants generated in this study. **(b)** SDS-PAGE of purified IgG1 WT and each mutant under reduction and non-reduction conditions. In the reduced SDS-PAGE, the upper and lower arrows represent the position of the heavy chain (around 50 kDa), and the light chain (around 25 kDa), respectively. In the gel under non-reducing conditions, most ΔP230 and essentially all CSCS were found to run at half molecular size (around 75 kDa) compared with the full length IgG1 (around 150kDa) as indicated. **(c)** SEC chromatograms of ΔP230 and CSCS mutants. The elution volume of CSCS was 12.3 ml (similar to WT) whereas ΔP230 eluted at two positions, with the large fraction at 13.7 and a minor fraction at 12.3 ml. **(d)** SDS-PAGE of IgG1-Fc WT (Asp221-Lys447) and ΔP230-Fc after purification. The gel was obtained under reducing and non-reducing conditions as indicated. The upper arrow represents the dimeric form (around 50 kDa) of the Fc fragment, and the lower arrow represents the monomeric form (around 25 kDa). **(e)** SEC chromatograms of WT-Fc and ΔP230-Fc. The position of the elution peak of WT Fc and ΔP230 were 9.1 ml and 9.2 ml, respectively.

Size exclusion chromatography (SEC) was conducted to further characterize the molecular weight of the ΔP230 and CSCS species in solution. The CSCS appeared as a single peak at 12.3 ml, showing a position similar to that of WT IgG1, and thus indicating that in solution CSCS adopts the characteristic fully assembled structure of WT IgG1 (150 kDa) (Fig. 1c). Meanwhile, ΔP230 appeared in two different peaks, a minor one at 12.3 ml similar to that of WT IgG1, and a major peak at 13.7 ml, roughly corresponding to the half-sized antibody (i.e. 75 kDa). This later result was unexpected, because the architecture of IgG1 in solution, even in the double mutant CSCS lacking disulfide bonds, is maintained by the interactions between the C_H_3 domains far from the hinge region. The deletion ΔP230, located in a region remote from the C_H_3 domain seems to exert a significant long-distance effect that compromised the native architecture of the IgG1 molecule.

To elucidate the contribution of the Fab region in the formation of half-IgG in ΔP230, we examined the effect of ΔP230 on the dimerization of the Fc fragment (Asp221 to Lys447). The position of the bands obtained by SDS-PAGE under reducing and non-reducing conditions for Fc of ΔP230 and for WT Fc were nearly indistinguishable (Fig 1d). In both cases, a single band at around 50 kDa corresponding to the dimeric form, and a band at around 25 kDa corresponding to the monomer were found under non-reducing and reducing conditions, respectively. The SEC profile of each construct was also very similar to each other: for the Fc fragment of ΔP230 the band was centered at 9.2 ml, whereas that of the Fc fragment of WT IgG1 was determined to be 9.1 ml (Fig. 1e). Collectively, these data have clearly established that the dimerization of the heavy chain of ΔP230 is dramatically reduced in the full-length antibody containing the Fab region, but not when only the Fc region is examined.

### Changes in the disulfide bond pattern in the hinge region caused by ΔP230

In WT IgG1, Cys226 and Cys229 residues of one heavy chain form intermolecular disulfide bonds with their counterparts, also Cys226 and Cys229, of the other heavy chain. In contrast, the data from above suggested that the formation of disulfide bonds in the deletion mutant ΔP230 was elusive, since the major fraction of this mutant in solution and SDS-PAGE under reductive conditions was the half IgG architecture. To investigate the state of these pair of cysteine residues (Cys226, Cys229) in a rigorous fashion in the deletion mutant ΔP230, disulfide quantification was performed by LC-MS/MS. The experiments were performed for the alkylated and denatured samples of half IgG fraction of ΔP230 and WT IgG1. After digestion with trypsin and LysC under non-reducing conditions to compare the detected peaks of trastuzumab WT and ΔP230 (Fig. 2). In the WT IgG1, as high as 97% of the molecule formed intermolecular disulfide bonds with the partner molecule, whereas in ΔP230, that percentage represented less than 2% of the molecules. This did not mean that the Cys residues were found in the free form, but instead 98% engaged with a partner Cys by intramolecular (intrachain) disulfide bonds (Table 1).

**Figure 2.**
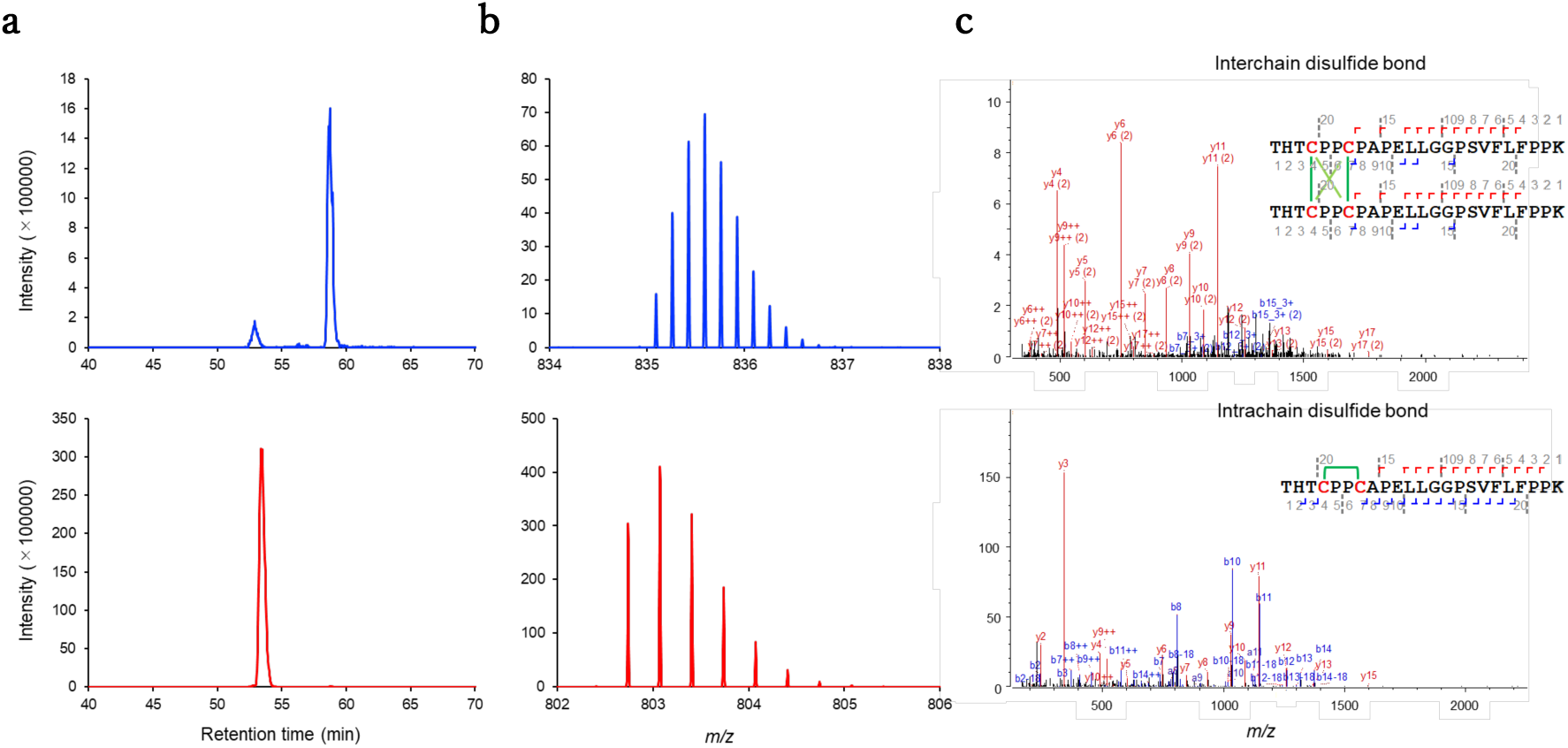
Analysis of hinge disulfide bond formation by LC-MS. **(a)** Extracted ion chromatograms of hinge peptides (WT: THTCPPCPAPELLGGPSVFLFPPK, ΔP230: THTCPPCAPELLGGPSVFLFPPK), with interchain disulfide bond in WT (m/z 835.0890 ± 0.01) (top) and T226–K248 with intrachain disulfide bond in ΔP230 (m/z 802.7380± 0.01) (bottom). **(b)** The mass spectra at 58.75 min in WT (top) and 53.40 min in ΔP230 (bottom). **(c)** The Higher-Energy Collisional Dissociation mass spectra of hinge peptides with two interchain disulfide bonds in WT (top) and intrachain disulfide bond in ΔP230 (bottom).

**Table.1.** Quantification of hinge disulfide bonds of WT and ΔP230 IgG1 determined by LC-MS/MS.

|  | Interchain S-S | Intrachain S-S |
| --- | --- | --- |
| WT | 99.2% | 0.8% |
| ΔP230 | 1.9% | 98.1% |

### Analysis of the structural effects of ΔP230 on the hinge and Fc by Nuclear Magnetic Resonance

To analyze the structural effect of the deletion of Pro230 on the hinge and Fc at the atomic level, we conducted NMR analysis of the Fc fragment. ^15^N labeled Fc was prepared with the Expi293 expression system, in which the medium was exchanged to contain amino acids labeled with ^15^N at the time of transfection. The yield of ^15^N labeled Fc yielded was nearly comparable to that of Fc expressed in normal medium (∼3 mg of pure protein was obtained from 25 ml of transfected cells). This expression method was, for the first time, explored and successfully applied for the purpose of obtaining a glycoprotein fully labelled with ^15^N from transient expression. This method will be useful for the mutational analysis of glycoproteins in general, where the establishment of stable expression systems are time consuming. We compared ^1^ H–^15^N HSQC spectra between WT Fc and ΔP230 Fc (Fig. 3a) and found structural differences in the flexible hinge region. We could selectively observe this region due to the high mobility. We confirmed that residues showing chemical shift differences are confined near the region of the hinge (from Cys226 to Gly237) (Fig. 3b).

**Figure 3.**
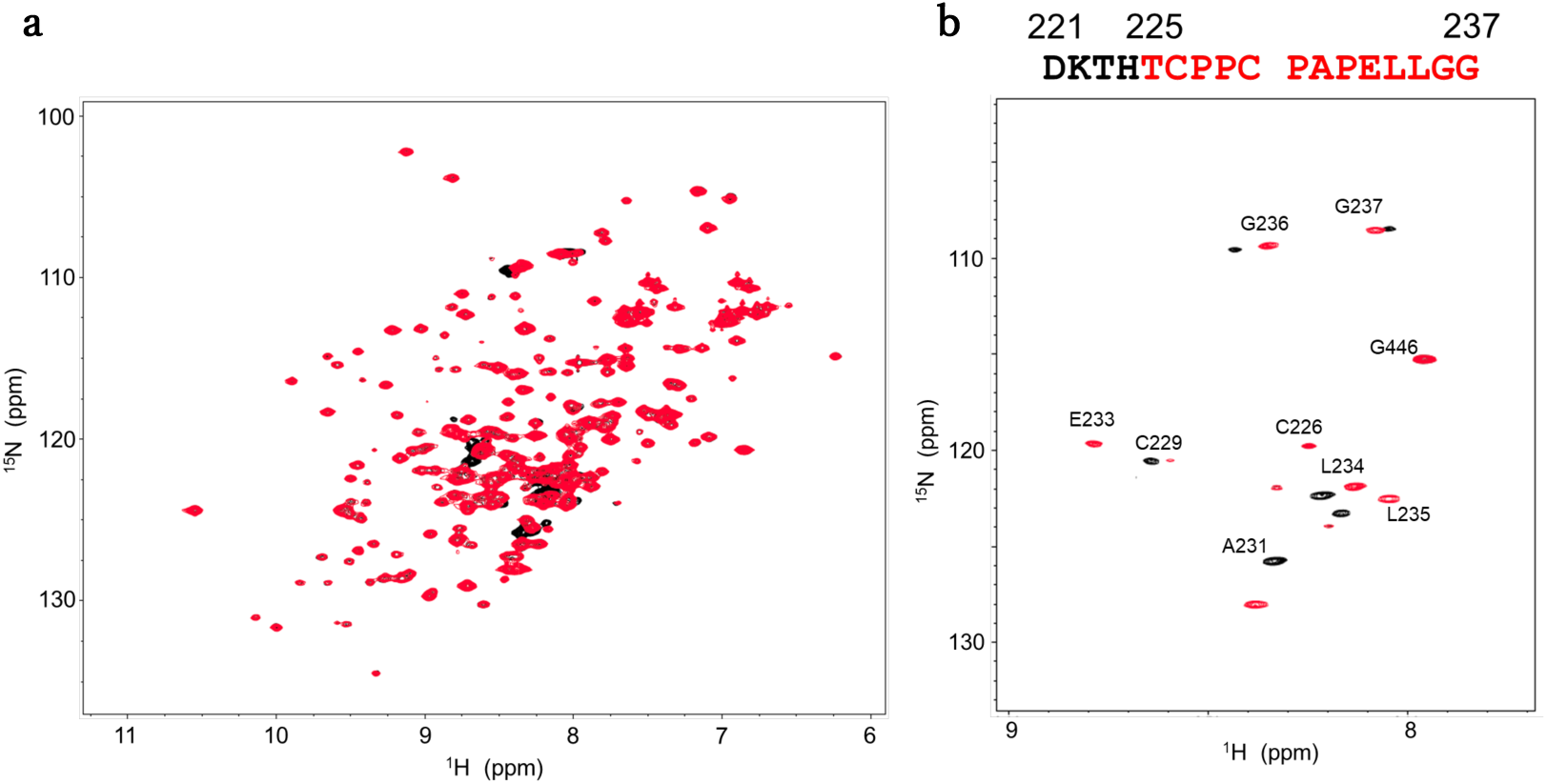
NMR analysis of the effect of ΔP230 mutation. Comparison of ^1^H-^15^N HSQC spectrum of **(a)** all NMR signal and **(b)** the NMR signal of the hinge region for the ^15^N-labeled IgG1-Fc WT and ΔP230. The WT and the ΔP230 Fc spectra are shown is in black and red, respectively.

### Evaluation of the interaction of hinge mutants of IgG1 with FcγRI

To investigate the effect of each mutation on the function of IgG1, the binding affinity to FcγRI was determined by the technique of surface plasmon resonance (SPR). FcγRI (WT) was immobilized on a CM5 surface chip by the amine coupling method (immobilized level ∼5,000 RU), whereas WT IgG1 or mutants were flowed as the analyte. The kinetic parameters, association rate constant *k*_on_ and dissociation rate constant *k*_off_, were obtained by a fitting procedure using the software BiaEvaluation (Fig. S1). The dissociation constant (*K*_D_) was determined from the kinetic parameters *k*_on_ and *k*_off_ for WT and each of the mutants (Table 2). The binding of WT to the receptor produced a sharp increase in the SPR signal, followed by a slow and steady decrease in signal after the antibody was ceased to be injected in the chip, resulting in a high affinity dissociation constant of 0.89 nM. When the analyte injected in the chip was the deletion mutant ΔP230, the association phase slowed down significantly, whereas the dissociation constant accelerated slightly, resulting in a still relatively strong binding affinity to FcγRI (*K*_D_ = 23.5 nM). Although this value is approximately 26-fold worse than WT IgG1, it is certainly still significant given that most of the antibody is now in the monomeric form. Other single deletion mutants (ΔA231 to ΔE233) targeted residues in the close vicinity to the binding interface of IgG1 with FcγRI^28^, and their deletion resulted in moderate reductions of affinity (2-to 4-fold lower affinity compared with WT). For the double Cys mutant CSCS, the resulting value of affinity was similar to that of the WT (*k*_on_ = 5.9 × 10^5^, *k*_off_ = 5.5 × 10^−4^, *K*_D_ = 0.92 nM), suggesting that the presence (or lack thereof) of disulfide bonds in the hinge region exerts a negligible effect on protein-protein interactions between IgG1 and its cellular receptor FcγRI.

**Table.2.** Kinetic parameters for IgG1-FcγRI interactions obtained by SPR measurements.

| | $k_{\text{on}}$ (1/Ms) | $k_{\text{off}}$ (1/s) | $K_{\text{D}}$ (nM) |
| --- | --- | --- | --- |
| WT | $5.5 \times 10^5$ | $4.9 \times 10^{-4}$ | 0.89 |
| ΔP230 | $4.3 \times 10^4$ | $1.0 \times 10^{-3}$ | 23.5 |
| ΔA231 | $2.5 \times 10^5$ | $5.8 \times 10^{-4}$ | 2.3 |
| ΔP232 | $6.2 \times 10^5$ | $1.0 \times 10^{-3}$ | 1.7 |
| ΔE233 | $5.0 \times 10^5$ | $1.7 \times 10^{-3}$ | 3.5 |
| CSCS | $5.9 \times 10^5$ | $5.5 \times 10^{-4}$ | 0.92 |

High-speed atomic force microscopy (HS-AFM) was subsequently employed to examine the interaction between antibody and receptor at the single-molecule level. HS-AFM images of IgG1 WT showed three bright spots in delineating a Y-shape consistent with the expected arrangement of a full-length antibody (Fig. 4a), whereas only two bright spots were seen in ΔP230 (Fig.4b), validating the observation above that this mutant exists mainly in a half-IgG architecture. FcγRI was immobilized using its C-terminal His_6_ tag on the mica surface, IgG1 WT or ΔP230 added in separate experiments, and their interaction was monitored (Fig.4c-e). Fig. 3c displays successive HS-AFM images of IgG1 WT. Here, the positions of IgG1 WT and FcγRI overlap, rendering them indistinguishable. However, since IgG1 WT is not stationary due to diffusion on surfaces lacking FcγRI, it is believed that the images depict IgG1 WT bound to FcγRI immobilized on the Ni2+/mica surface. This observation suggests that IgG1 WT and FcγRI interact with a 1:1 stoichiometry. On the other hand, when ΔP230 was in solution, two distinct cases were observed: a single ΔP230 bound to FcγRI (Fig. 4d) and two ΔP230 bound to FcγRI (Fig. 4e), indicating that the interaction stoichiometry ΔP230-to-FcγRI fluctuated between 1:1 (mol:mol) and 2:1 (mol:mol). Since in previous crystal structures, it is shown that FcγRI and IgG1 interact in a 1:1 binding mode with two chains of Fc both contributing to the interaction, it is reasonable to see that two chains (one from each ΔP230 half IgG) assemble together in the presence of FcγRI. In addition, the (1:1) stoichiometry corresponding to one half-chain antibody and one molecule of the receptor is also observed, consistent with the asymmetric interaction surface described in the crystal structure. In the structure, it was suggested the possibility that a single Fc chain could bind to this receptor given the much larger interaction surface of one of the Fc with respect to the second one.^28,29^

**Figure 4.**
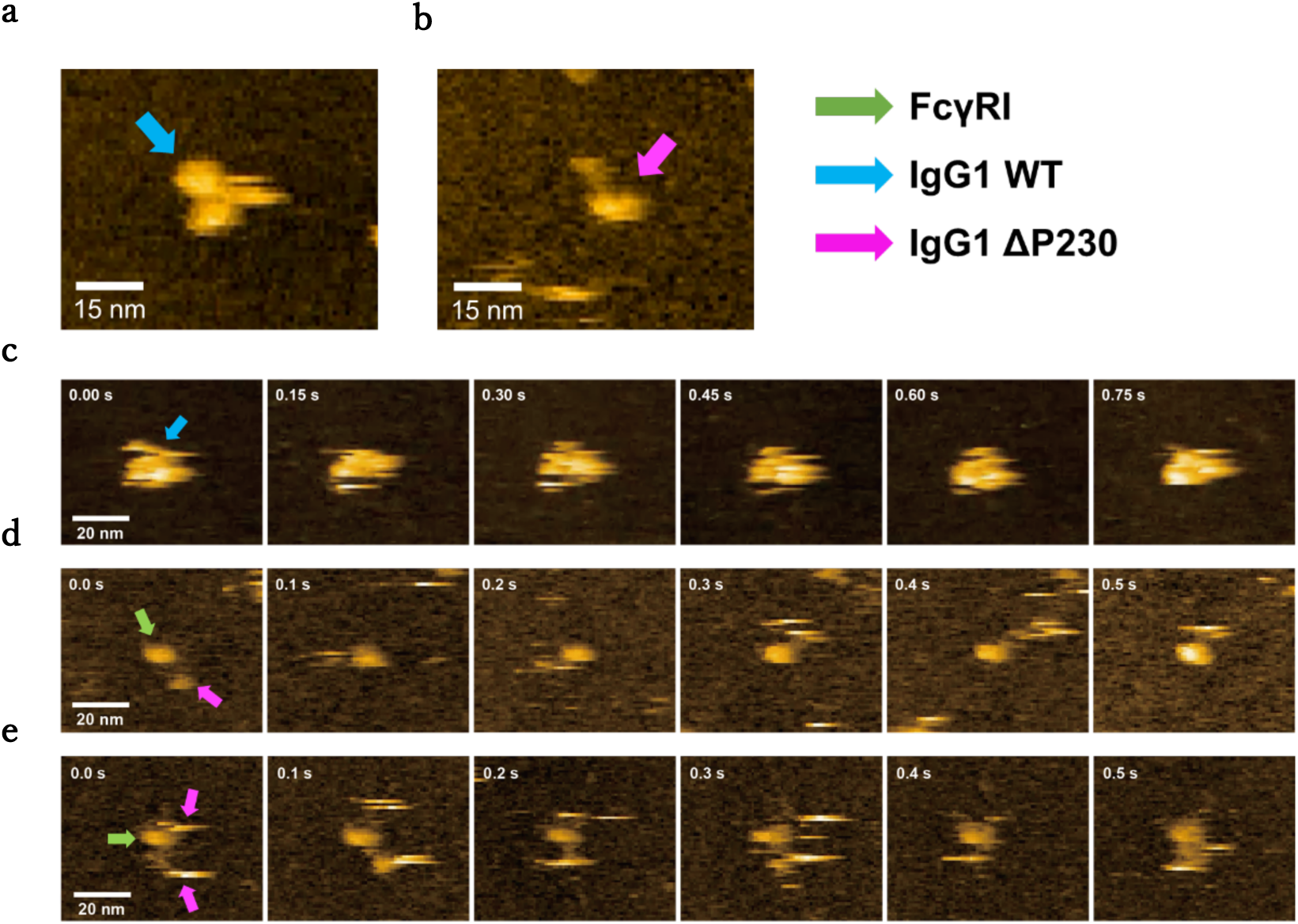
HS-AFM observation of the interaction of WT and ΔP230 IgG1 with FcγRI. Representative snapshots of **(a)** WT IgG1 and (b) ΔP230 captured by HS-AFM. Successive HS-AFM images of **(c)** WT IgG1 and **(d), (e)** ΔP230 bound to FcγRI immobilized on Ni^2+/^mica surface. Images (a) were taken at an imaging rate of 0.1 5 s/frame, (d) and (e) at 0.1 s/frame. WT IgG1 is indicated by green arrows, ΔP230 by magenta arrows, and FcγRI by green arrows. While WT formed only 1:1 stoichiometric complexes with FcγRI, ΔP230 formed not only 1:1 but also 1:2 stoichiometric complexes with FcγRI.

### Examination of effector-activity of ΔP230 via FcγRI

Since the data above indicated that the interaction of ΔP230 and FcγRI are maintained, we examined the physiological relevance of this interaction by analyzing their activity at the cellular level (Fig. 5). In particular, we examined various effector activities derived from the interaction with various specific Fcγ receptors. The results of activity measurements showed that ΔP230 maintains the effector-activity via FcγRI although it showed reduced activity. We confirmed that ΔP230 does not have FcγRIIa or FcγRIIIa mediated activity, even when subjected to higher concentrations of antibody compared to FcγRI.

**Figure 5.**
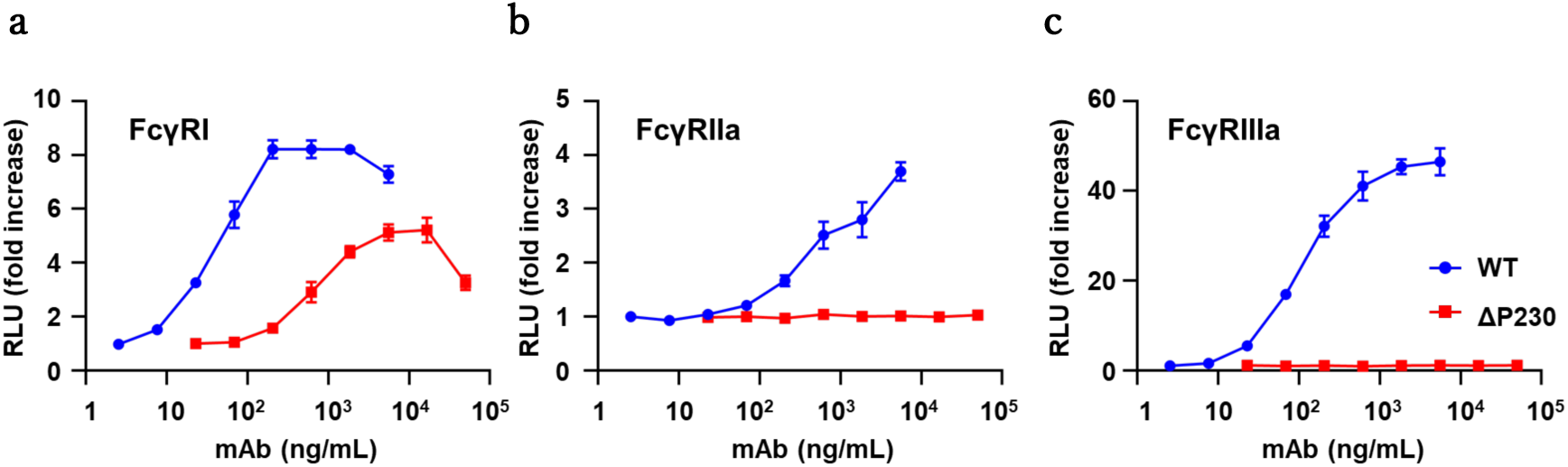
Effector function of WT IgG1 and ΔP230. The effector function of antibodies were evaluated with three different reporter assays based on the activities of **(a)** FcγRI, **(b)** FcγRIIa, and **(c)** FcγRIIIa. WT and ΔP230 are represented by blue and red, respectively.

**Figure 6.**
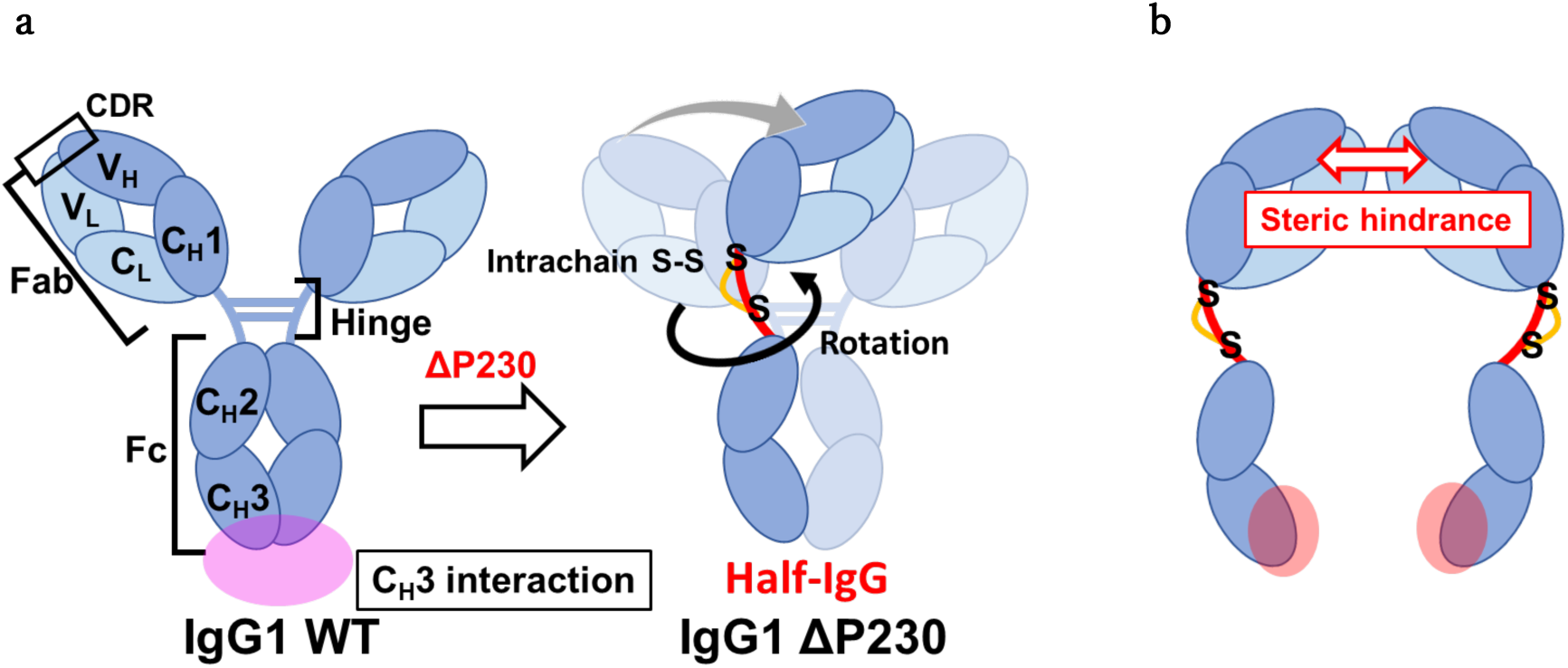
Proposed mechanism of half IgG formation in ΔP230. **(a)** ΔP230 causes disturbance in the surrounding residues, leading to a change in the disulfide bond pattern in the lower hinge from an intermolecular to an intramolecular arrangement. The strong energy of the disulfide bonds (a type of covalent bond) is sufficient to change the relative orientation between the Fab and Fc regions. **(b)** The new orientation of the large Fc and Fab regions force the IgG1 molecule to split as half IgG. A possible reason is that the strain between Fab and Fc regions disrupt the stabilizing interaction between C_H_3 domains, possible by steric hindrance imposed by the Fab regions. Light and heavy chains are represented by light blue and darker blue, respectively.

## Discussion

In previous studies the effect of hinge modification on various properties of IgG1 were extensively conducted for the core and upper hinge region in previous reports.^5–14^ To further establish the role of the hinge region on the architecture and the function of IgG1, we prepared systematic deletion of residues in the lower hinge region. We show that the deletion ΔP230 (located between the hinge cysteine residues) results in a predominant population of half IgG molecules in solution in a ratio monomer:dimer of approximately 6:1. This conclusion is based not only on data from SDS-PAGE under non-reducing conditions, but also in chromatographic analysis by SEC and HS-AFM images. In the previous research, the structural integrity of IgG1 mutants were investigated only by SDS-PAGE. Therefore, the previous knowledge of the hinge mutants is that some mutants (ΔP227ΔP228, ΔT224ΔH225ΔP227ΔP228, CPGGGPCP and P227W P228W) are defected in interchain disulfide bond formation.^10^ In light of our findings that ΔPro230 indeed exists as a half IgG molecule in solution, it is possible that some of the previously reported hinge mutants which do not form interchain disulfide bonds might also exist as half IgG entities.

In the investigation of the mechanism of half IgG formation, we found that ΔP230 alters the disulfide bond pattern in the hinge, changing from a predominantly intermolecular arrangement (WT form) to largely intramolecular pattern in the mutant ΔP230 (Fig. S1). In addition, the Fab region decisively contribute to the formation of the half IgG (Fig. 1d) and comparative NMR data determined the structural differences between the Fc region of WT and ΔP230 mutant are restricted to the hinge region. Collectively, these data lead us to speculate that the alteration in the disulfide patter in the lower hinge upon elimination of Pro230 results in a the spatial rearrangement between Fab and Fc, with potential clashes between the two Fab regions and potential long-distance effect on the dimerization surface in the C_H_3 domains as shown in Figure 5.^30–32^

Importantly, the half IgG molecules retain the ability to bind to the FcγRI as determined by HS-AFM with different stoichiometric ratios (1:1 and 2:1). At the cellular level, this translates in a diminished but still significant effector activity of half IgG mediated by FcγRI. No effector activity linked to FcγRIIa or FcγRIIIa was observed. Previous studies have reported complex crystal structures of FcγRI and Fc fragments^28,29^, indicating that these two molecules interact through two different interfaces. Since the half IgG entity has only one interface, there are two possible binding modes observed, suggesting that it is active only with respect to FcγRI, a high-affinity Fcγ receptor subtype that can retain binding through only one of the interfaces

In summary, our study brings new insight on the effect of the lower hinge region of IgG1 on the formation of the architecture (higher-order structure) of IgG1 and its influence in the activity of the antibody at the cellular level. The half IgG molecule found in this study is a useful tool to understand the role of FcγRI by its specific activity against FcγRI, and further research could result in a novel antibody drug modality.

## Materials & Methods

### Cells

SK-BR-3 cells (an HER2^+^ human breast cancer cell line; ATCC, HTB-30) was obtained from ATCC. Jurkat/FcγR/NFAT-Luc cells, which express human FcγRI, FcγRIIa, or FcγRIIIa and NFAT-driven luciferase reporter, were developed previously.^33,34^ The cells were maintained at 37 °C in 5% CO2 with RPMI1640 medium (Thermo Fisher Scientific, Waltham, MA) supplemented with 10% heat-inactivated fetal bovine serum.

### Sample Preparation

The mammalian cell expression system Expi293 was used to express anti-HER2 humanized IgG1 Trastuzumab WT and mutants. Plasmids of Trastuzumab WT and mutants in pcDNA3.4 vector were transfected into Expi293 cells, a mammalian cell expression system. The Gibco™ Expi293™ Expression System Kit (Thermo Fisher Scientific) and Gxpress 293 Transfection Kit (GMEP, Fukuoka, Japan) were used for protein expression. Transfection and culture condition were performed according to the kit protocol. Uniform ^15^N labeling of the IgG1-Fc glycoproteins was achieved by cultivating the transfected cells in medium containing the ^15^N-labeled amino acid mixture.^35^ The components of medium are listed in TableS1. Four days after transfection, collected cells were separated by centrifugation and the supernatant was subjected to affinity purification using Hitrap™ ProteinA HP Columns (Cytiva, Tokyo, Japan) which was equilibrated with 20 mM phosphate buffer pH7. After washing column by equilibration buffer, Trastuzumab was eluted by 100 mM citric buffer pH 2.7 and neutralized by 1 M Tris-HCl pH 9. Size exclusion chromatography purification (SEC) of Trastuzumab WT and mutants was performed using superdex 200 increase 10/300 GL (Cytiva) equilibrated with PBS, and SEC of Trastuzumab WT Fc and ΔP230 Fc was performed using superdex 75 increase 10/300 GL (Cytiva) equilibrated with PBS.

Galactosidase treatment of Fc fragments for NMR measurements was performed using buffer containing 50 mM acetic acid pH 5, 10 mM MnCl_2_. The N-glycans of Fc were unified by adding 1 nmol of β1-4 Galactosidase S (New England Biolabs, Tokyo, Japan) per 1 unit of glycan terminus (mixed so that the enzyme solution was 10% of the total reaction solution) and reacting at 37°C for 72 hours.^36^ After the reaction, affinity purification was performed by ProteinA column to remove the enzyme. Human FcγRⅠ was purchased from Acro Biosystems, Co., Ltd.

### HS-AFM

The HS-AFM observations were performed with tapping mode at room temperature. A microcantilever (AC7, Olympus Corporation, Tokyo, Japan), with a resonance frequency of approximately 600 kHz in solution, a spring constant of about 0.2 N/m, and a *Q* value of approximately 0.7, was used. The cantilever was oscillated with a free oscillation amplitude of approximately 1 nm. During imaging, the distance between the probe and the sample was maintained by feedback control to keep the oscilation amplitude between 0.6 and 0.8 nm. To examine the interaction between FcγRⅠ WT and IgG1, FcγRI WT with a His tag at the C-terminus was immobilized on a Ni^2+^-coated mica Initially, 2 mM NiCl2 was deposited on a freshly cleaved mica surface and incubated for 3 minutes, followed by washing with MlliQ-water. Afterward, 4 μl of FcγRI WT was added and allowed to incubate for 5 minutes. Unbound FcγRI WT was then washed away with an observation buffer (50 mM Tris-HCl, 100 mM NaCl, pH 8). Subsequently the HS-AFM imaging was carried out in the same buffer. During AFM observation, IgG1 was added to the observation buffer and its interaction with FcγRI WT immobilized on the substrate was observed. Here, IgG1 rapidly diffuses on the Ni^2+^/mica surface in the presence of 100 mM NaCl, rendering it unobservable by HS-AFM. Consequently, when IgG1 was visualized on the mica surface, as shown in Fig. 4a, the HS-AFM imaging was carried out in the observation buffer without NaCl.

### Surface Plasmon Resonance

Biacore8K (GE Healthcare, Chicago, IL) was used for interaction evaluation using SPR. FcγRⅠ WT was diluted in acetic acid buffer pH 5 and immobilized on the surface of Series Sensor Chip CM5 (GE Healthcare) by amine coupling method. (Immobilization level was ∼5000 RU) To reduce bulk effects, Trastuzumab sample were dialyzed against PBS + 0.005% Tween 20 buffer. After dialysis, concentrations of Trastuzumab were adjusted using the dialysis outer solution. PBS + 0.005% Tween20 was used for running buffer with the flow rate of 30 µl/min. Contact time of the analyte was 180 seconds, and dissociation time was 600 seconds. Regeneration was performed in 50 sec using 1M arginine pH 3.1. Fitting was performed on the resulting sensor gram using Biacore^TM^ Insight Evaluation Software.

### NMR

The Fc fragments were dissolved at a concentration of 6 mg/ml in 0.25 ml of 5 mM sodium phosphate buffer (pH 6.0) containing 50 mM NaCl and 10% (v/v) D_2_O. NMR spectra were acquired at 42°C using an AVANCE NEO 800 (Bruker BioSpin, Billerica, MA) spectrometer. Chemical shifts of ^1^H were referenced to DSS (0 ppm), and ^15^N chemical shifts were referenced indirectly using the gyromagnetic ratios of ^15^N and ^1^H (γ^15^N/γ^1^H = 0.10132905). All NMR data were processed using topspin software and further analyzed using NMRview.

### LC-MS/MS

Sample IgGs were incubated with 5 mM Iodoacetamide 50 mM Ammonium bicarbonate at 25°C for 1 hour for alkylation of the free cysteine residues that might be present, and then guanidine hydrochloride was added at a final concentration of 6 M for 1 hour at 25°C for denaturation. After desalting and buffer-exchanged to 50 mM Ammonium bicarbonate with Zeba Spin Desalting Columns 40K, 0.5 mL (Thermo Fisher Scientific), LysC: trypsin: IgG1 was added at a ratio of 0.33:1:20, incubated at 37°C for 6 hours, and the reaction was stopped by formic acid at 1% final concentration. LC treatment was performed using the Ultimate 3000 (Thermo Fisher Scientific). The PepMap100 C18 (5 μm 0.3 x 5 mm, Thermo Fisher Scientific) and the ACQUITY BEH C18 (1.7 µm 1.0 × 100 mm, Waters) were used for desalination and separation. LC conditions used for separation were 45 μL/min with a 60 min gradient of 2%–35% with 0.1% formic acid in water to 0.1% formic acid in acetonitrile. Mass spectrometric analyses were performed using a Q-Exactive HFX (Thermo Fisher Scientific). The resolution was 120,000 for full MS scan, the mass range (m/z) was 300–2000. A subsequent ddMS/MS scan was performed by higher energy collisional dissociation with collision energies 27% and resolution 30,000. The data were analyzed using Byos v5.4.52 (Protein Metrics, Boston, MA).

### FcγR reporter cell assay

FcγR effector activity by IgG1 molecule was confirmed by FcγR reporter cell assay. SK-BR-3 cells were seeded at 5 × 10^3^cells/well in 96 well plate and incubated at 37°C for 24 hours, then the medium was removed and washed with PBS. FcγR reporter cells suspended in Opti-MEM I Reduced Serum Medium (Thermo Fisher Scientific) were then added at 1×10^5^ cells/well, and WT trastuzumab and ΔP230 were added at respective concentrations (WT 2.5 ng/mL∼5.6 µg/mL, ΔP230 22.9 ng/mL ∼ 50.0 µg/mL) and incubated at 37°C for 5 hours, followed by luciferase assay. The luciferase activities were measured by using a ONE-Glo Luciferase Assay System (Promega, Madison, WI) and an Ensight Multimode Plate Reader (PerkinElmer, Waltham, MA) according to the manufacturer’s protocol.

## Acknowledgements

We thank Yukiko Isono (Institute for Molecular Science) for her help in the preparation of the protein samples for NMR.

---

**Figure S1.**
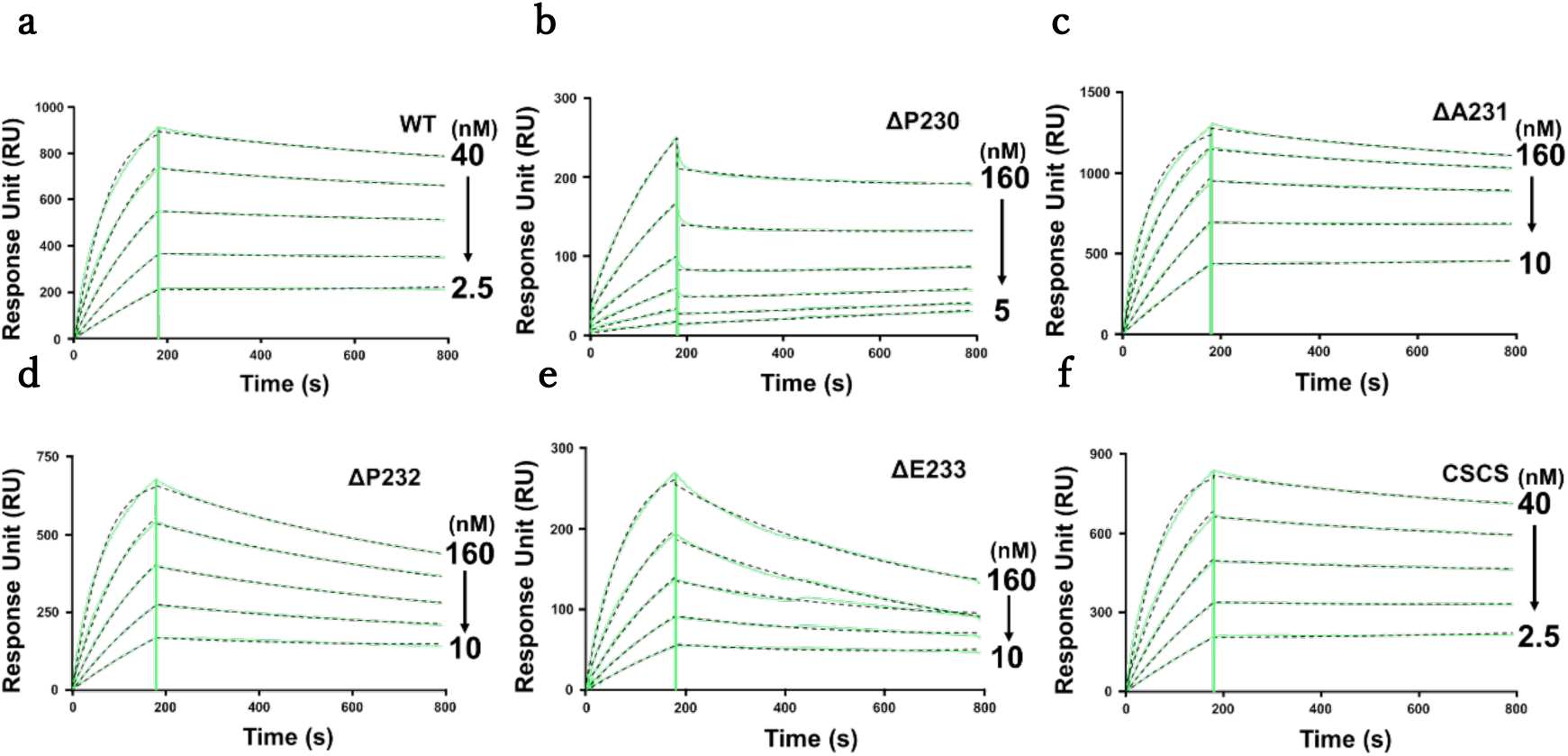
Sensor gram of IgG1-FcγRI interactions obtained by SPR measurements. Sensorgrams for IgG1 (a) WT and (b-f) each mutants against FcγRI obtained by SPR measurements. The concentrations of the analyte molecules were measured by double dilution from the highest to the lowest concentrations.

**Table S1.**
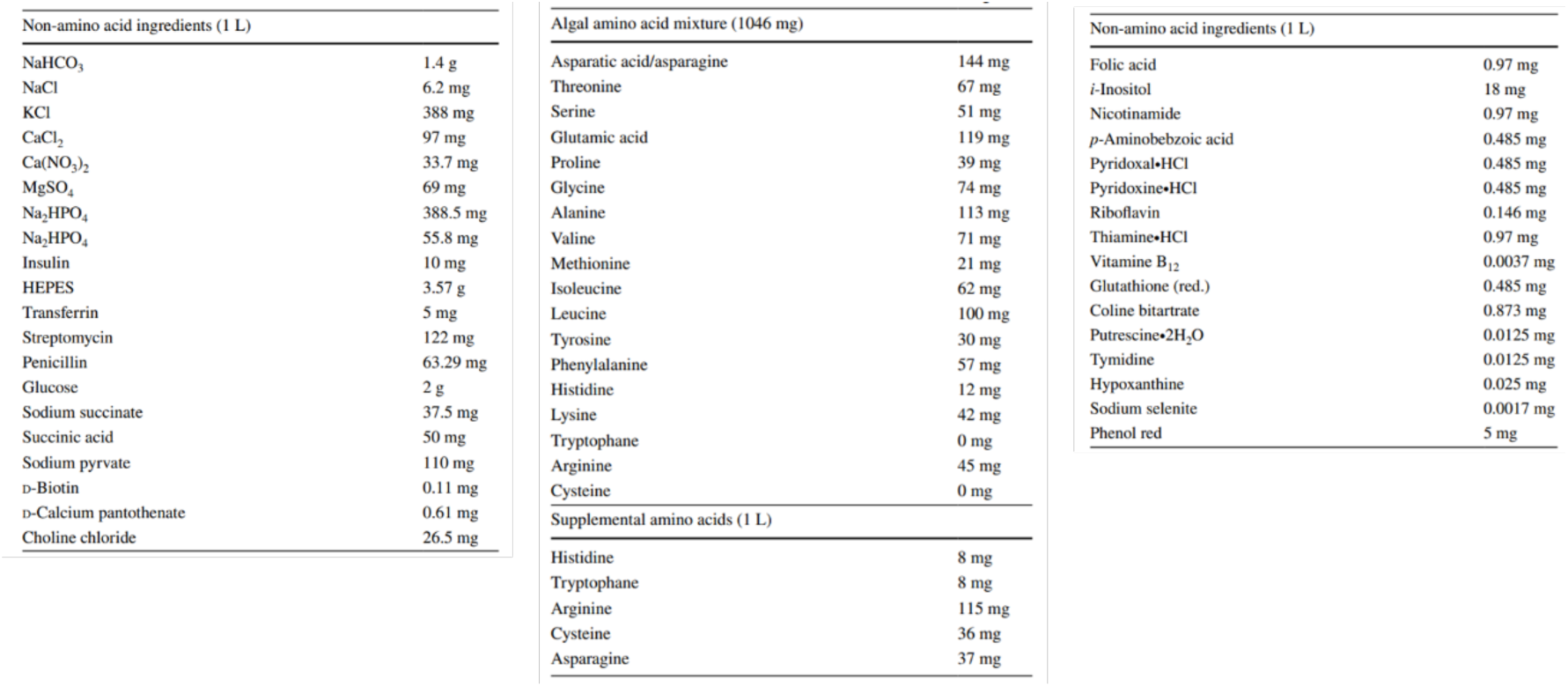
Composition of ^15^N-labeled NYSF medium.

**Movie S1 | HS-AFM movie of WT IgG1 bound to FcγRI immobilized on a mica surface.**

The typical Y-shaped structure of WT IgG1 is visible, fluctuating in solution. Unbound WT IgG1 molecules are observed diffusing on the mica surface after 10 seconds in the movie. The imaging rate was 0.15 s/frame.

**Movie S2 | HS-AFM movie capturing a single monovalent ΔP230 IgG1 binding to FcγRI immobilized on a mica surface.**

The relatively stationary bright spots in the center correspond to the Fab region of monovalent IgG1 bound to FcγRI, while the highly fluctuating region is considered to be the Fc region of IgG1. The imaging rate was 0.1 s/frame.

**Movie S3 | HS-AFM movie capturing dual ΔP230 IgGs binding to FcγRI immobilized on a mica surface.**

The imaging rate was 0.1 s/frame.

